# Multifaceted proteome analysis at solubility, redox, and expression dimensions for target identification

**DOI:** 10.1101/2023.08.31.555796

**Authors:** Amir Ata Saei, Albin Lundin, Hezheng Lyu, Hassan Gharibi, Huqiao Luo, Jaakko Teppo, Xuepei Zhang, Massimiliano Gaetani, Ákos Végvári, Rikard Holmdahl, Steven P. Gygi, Roman A. Zubarev

## Abstract

Multifaceted interrogation of the proteome deepens the system-wide understanding of biological systems; however, mapping the redox changes in the proteome has so far been significantly more challenging than expression and solubility/stability analyses. Here, we devise the first high-throughput redox proteomics approach integrated with expression analysis (REX) and combine it with Proteome Integral Solubility Alteration (PISA) assay. The whole PISA-REX experiment with up to four biological replicates can be multiplexed into a single tandem mass tag TMTpro set. For benchmarking this compact tool, we analyzed HCT116 cells treated with auranofin, showing great improvement compared with previous such studies. Then we applied PISA-REX to study proteome remodeling upon stimulation of human monocytes by interferon α. We also studied the proteome changes in plasmacytoid dendritic cells isolated from wild type vs. *Ncf1*- mutant mice treated with interferon α, showing that NCF1 deficiency enhances the STAT1 pathway and modulates the expression, solubility and redox state of interferon-induced proteins. Providing comprehensive multifaceted information on the proteome, the compact PISA-REX has the potential to become an industry standard in proteomics and to open new windows into the biology of health and disease.

## Introduction

Chemical proteomics employs tools for interrogation of biological systems and their interactions with small molecules and other entities. Such tools can be used in target deconvolution and exploration of drug mechanisms of action by the analysis of changes in the protein abundance, solubility/stability and redox state upon treatment with a biological or chemical entity ^1^. Such a multifaceted proteome interrogation can be extremely resource demanding. To alleviate this problem we have recently developed a high-throughput version of Thermal Proteome Profiling (TPP) or MS-CETSA ^2,3^ called Proteome Integral Solubility Alteration (PISA) assay ^4^. Thereafter we combined PISA assay with expression proteomics (PISA-Express) in the same tandem mass tag (TMT) multiplexing set to simultaneously track protein solubility/stability and expression changes upon cellular transitions to and from pluripotency ^5^. Multiplexing the whole experiment, including the desired number of biological replicates (≥3), into a single TMT set is highly beneficial, as it practically eliminates the problem with missing values ^5^. With the emergence of higher multiplexing power of TMTpro-16 and - 18plex sets ^6,7^, it was tempting to integrate redox proteomics in the arsenal of these chemical proteomics tools for three-facet (3f) analysis of the proteome within the same multiplexed set. Indeed, redox proteomics, although not a recent technique ^8,9^, is rapidly gaining attention of the proteomics community ^10,11^.

Reactive oxygen species (ROS) are metabolic or signaling molecules that exert their effects by oxidizing protein amino acid residues, particularly cysteines (Cys) ^12^, which can lead to modulation of protein activity or dysfunction ^13,14^. Unbiased proteome-wide mapping of redox-regulated Cys residues, i.e. redox proteomics analysis, is therefore essential for understanding ROS-mediated events. Advances in mass spectrometry-based proteomics have brought about several strategies in this area, but the coverage of cysteinome remained greatly inferior compared to expression proteomics or protein solubility/stability analyses (i.e., hundreds of analyzed proteins versus >5000 proteins in a typical case). With Cys constituting 2.3% of the proteome’s amino acids ^15^, ≈20% of all tryptic peptides contain at least one Cys; therefore, redox proteomics has significant unrealized potential in terms of proteome coverage.

One of the pitfalls of redox proteomics is that Cys residues, besides being in a free thiol or disulfide state, may undergo several modifications, such as sulfenylation, sulfinylation, sulfonylation, nitrosylation and glutathionylation ^16^, which complicates peptide detection by database search. Furthermore, in most tissues and organelles, only around 10% of Cys residues are oxidized on average ^17^, and the low abundance often precludes the oxidized peptide to be identified, and even more so, quantified in an LC-MS/MS experiment. Since most redox proteomics techniques are based upon simultaneous quantification of both reduced and oxidized version of a given Cys-containing peptide to enable calculation of the oxidation occupancy ^1^, these techniques have achieved a rather low coverage. Extensive bioinformatics (e.g., imputing missing values), reverse labeling strategies with light and heavy isotopes and retention time matching provide only partial remedy ^18^.

Multiple strategies have been applied to enrich for Cys-containing peptides using isotope-coded affinity tags (ICAT) ^19^, OxICAT ^20^, irreversible isobaric iodoacetyl tandem mass tags (iodoTMT) ^21^, or OxiTMT ^22^, isotopic tandem orthogonal proteolysis-activity-based protein profiling (isoTOP-ABPP) ^23^, click chemistry ^24^, and Cys-reactive phosphate tags coupled to IMAC enrichment ^17^. While some of these approaches provide quite decent cysteinome coverage (≥1000 proteins per analysis), the enrichment step involves labor-intensive workflows with multiple washings that can result in sample loss and/or induction of spontaneous oxidation events. Importantly, upon Cys-peptide enrichment, protein abundance information is lost, which necessitates performing expression analysis as a separate experiment ^25^.

In a recent technique called Stable Isotope Cys Labeling with IodoAcetamide or SICyLIA ^25^, some of these issues have been addressed. In this technique, proteins are extracted in the presence of either light iodoacetamide (IAA) or heavy isotope labeled IAA to alkylate free Cys thiols in proteins by a carbamidomethyl group. After mixing equal amounts of modified light and heavy protein extracts representing two different samples under comparison, dithiothreitol (DTT) is used to reduce the oxidized thiols, which are subsequently alkylated with N-ethylmaleimide (NEM). The ratio of the heavy to light labeled Cys serves as a proxy for comparing the levels of reduced Cys in the two samples. This technique circumvents the need to measure the oxidized protein thiols (NEM-labeled peptides are simply ignored in SICyLIA) and therefore, using peptide fractionation prior to LC-MS/MS, one can achieve a satisfactory cysteinome coverage without any enrichment. While this approach provided by far the best redox proteome coverage (9,479 peptides in mouse cells and 4,415 in tissues, corresponding to 3,563 and 2,168 proteins, respectively, after keeping unique peptides quantified in 3 out of the 4 replicates), under dynamic cellular conditions a separate expression proteome analysis must be performed in parallel to SICyLIA in order to account for changes in the protein abundances. In the cited work, this was achieved by dimethylating tryptic peptides using light or heavy formaldehyde/sodium cyanoborohydride. The two data sets were then matched, and the redox proteomics data were normalized by the abundances of the corresponding proteins.

Since the redox and expression samples in SICyLIA undergo different processing and labeling procedures, combining their results into a single dataset can be challenging and may entail some data loss. Moreover, unless for each tryptic peptide both light and heavy dimethylated variants are quantified, missing values inevitably emerge. Another significant limitation is that only two samples can be multiplexed in redox or abundance analysis by dimethylation. In addition, heavy IAA is three orders of magnitude more expensive than light IAA (1 g of heavy IAA costs more than 1 kg of light IAA).

Having learned from the above techniques and attempting to reach the goal of multiplexing the whole experiment into a single TMT set, we address here the mentioned problems by introducing the REX proteomics approach that integrates the redox and expression facets. We also combine REX with PISA and remove the redundancy in expression analysis. To test this compact PISA-REX technique, we first applied it to HCT116 cells treated with the redox-active drug auranofin, which we have extensively characterized previously with expression, solubility and redox proteomics ^1^. Subsequently, we performed PISA-REX investigation of interferon α (IFN-α) stimulation in human monocytes, generating a resource for the community. Finally, we investigated proteome changes in plasmacytoid dendritic cells (pDCs) isolated from wild type vs. *Ncf1*-mutant mice treated with IFN-α, to uncover the key molecular events associated with this mutation in lupus. In all these cases, the treatments profoundly altered the redox state of the cellular proteome. With PISA-REX we identified the targets and the mechanisms of treatments and then validated the most important findings using orthogonal approaches.

## Results

### REX with separate redox and expression TMT channels

First, we combined expression proteomics with redox proteomics using separate TMT labels (channels). For expression analysis, the whole protein lysate was reduced with DTT and then labeled with IAA, as in conventional proteomics (**Fig. 1a**). For redox analysis, the normally dominant reduced Cys thiols were labeled with conventional IAA, after which oxidized Cys residues were reduced by DTT and blocked with NEM (**Fig. 1a**). The main purpose of NEM labeling was to stabilize the newly formed thiols and prevent secondary reactions; as in SICyLIA ^25^, no detection of the NEM-labeled peptides is required. Note that, as no isotope labeling is used in REX, the cost of sample preparation remains minimal. The proteins were then digested with trypsin, and tryptic peptides corresponding to control and treated samples were labeled with unique TMT labels ^7^. A PISA analysis was then performed as described earlier ^4,26^. All REX and PISA samples (treated vs untreated cells in three replicates) were then combined within a single TMTpro-18plex set, providing a 3-faceted (3f) PISA-REX experiment (**Fig. 1b**).

**Fig. 1.**
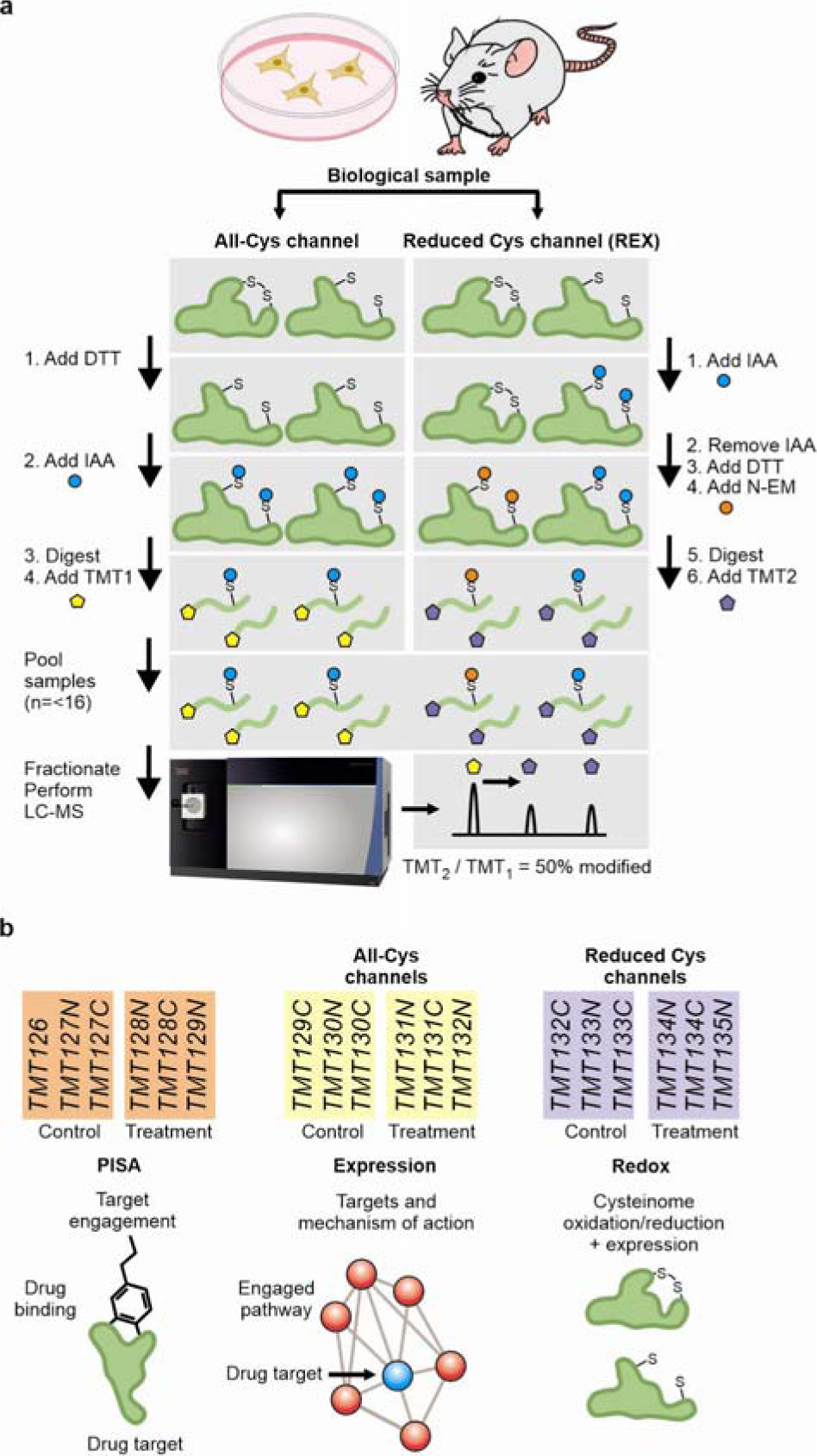
PISA-REX workflow and experiment design. **a,** The REX sample preparation protocol for one condition is shown. A given biological sample is divided into two portions. One portion intended for expression analysis is processed according to the routine proteomics workflow, with full reduction of Cys with DTT and alkylation with IAA. The other portion devoted to redox analysis is first labeled with IAA, and after removal of IAA, DTT is added to reduce the oxidized Cys and block them with NEM. **b,** The experimental design for inclusion of PISA and REX proteomics samples within a TMTpro-18plex set-up. The type of information obtained from each type of analysis is shown.

As proof of principle, we applied PISA-REX to HCT116 cells treated with auranofin, a redox active compound approved for treatment of rheumatoid arthritis. Auranofin is being repurposed in several clinical trials on ovarian cancer, lung cancer and chronic lymphocytic leukemia (trial numbers NCT01419691, NCT01737502, NCT01747798, and NCT03456700; see www.clinicaltrials.gov). Auranofin is an inhibitor of thioredoxin reductase TXNRD1 ^27^. We have recently applied comprehensive chemical proteomics tools to study auranofin, including TPP, Functional Identification of Target by Expression Proteomics or FITExP (based on expression proteomics) ^28^ and redox proteomics through sequential iodoTMT labeling ^1^. Therefore, to benchmark PISA-REX (dataset 1), we replicated the same experimental conditions, i.e. for PISA and redox analysis, cells were treated with 3 μM auranofin for 2 h, while for the expression assessment, cells were treated for 48 h at IC50 concentration (1.5 μM) (**Supplementary data 1** and **2** provide protein and peptide level data, respectively).

In our previous work the redox analysis covered only 2,129 peptides belonging to 1,383 proteins ^1^. In contrast, here (dataset 1) we quantified 98,945 peptides. Of the 18,538 Cys containing peptides, 17,089 carbamidomethylated molecules (13,907 unique sequences) were detected belonging to 5,029 proteins (92.2% IAA labeling efficiency).

Data processing was performed as follows. After removing the few peptides with missing values in TMT channels, peptide abundances were calculated by normalizing the intensities of the corresponding reporter ions by the total intensity of that TMT channel. For the PISA and expression dimensions, protein abundances were calculated as the summed abundances of all peptides belonging to a given protein (as given by Proteome Discoverer), and the fold changes in auranofin treatment vs control were calculated. Redox analysis was performed for carbamidomethylated peptides, the abundances of which were normalized first by the total intensity of that TMT channel and then by the summed abundance of non-Cys containing peptides from the same protein to adjust for the eventual protein abundance changes during treatment. Only peptides carrying the carbamidomethyl were considered. Finally, the log2 values of the fold-changes (FCs) of thus obtained values in auranofin treatment vs. control were calculated (**Fig. 2a**).

**Fig. 2.**
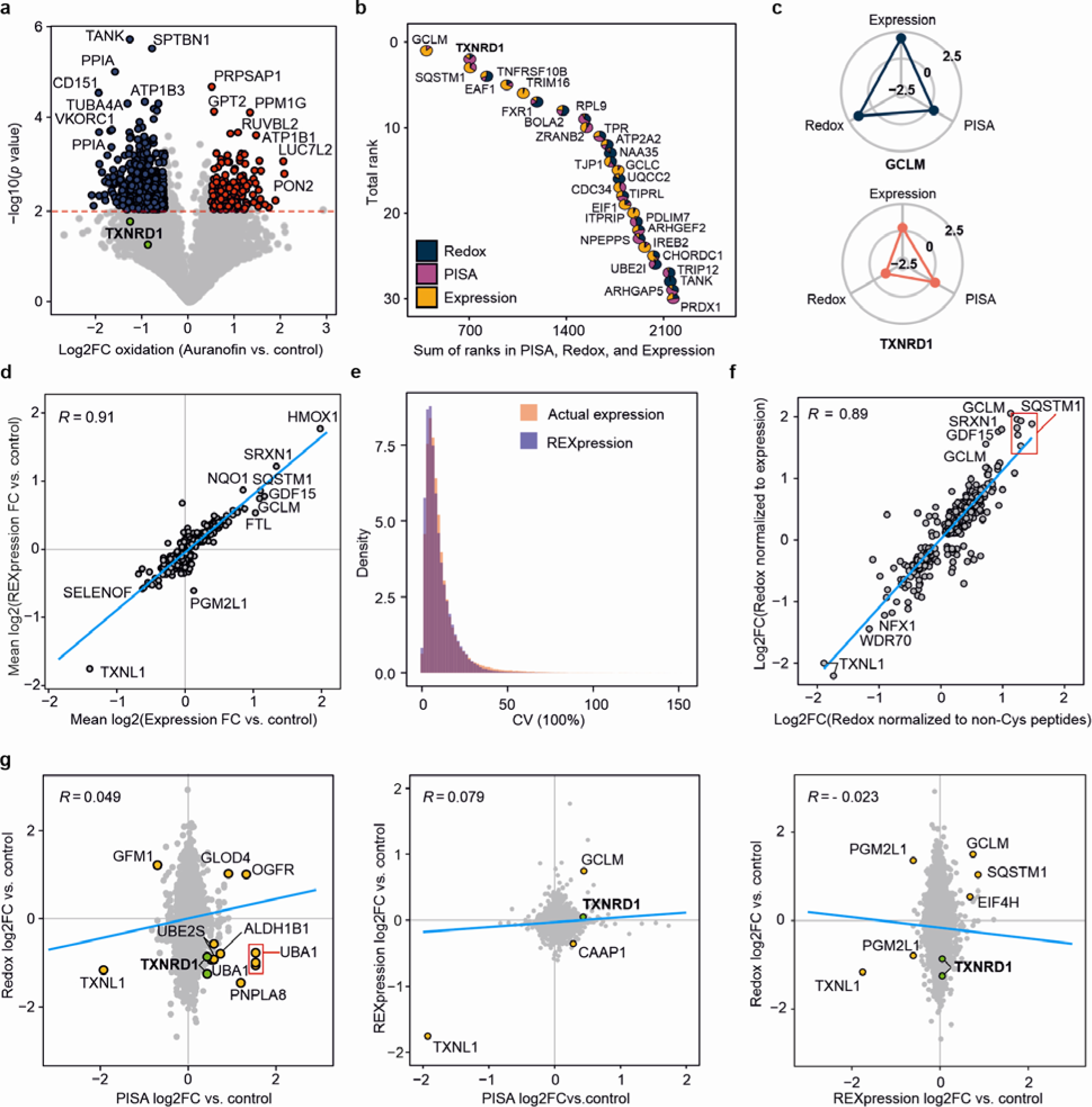
Benchmarking PISA-REX. **a,** A 2 h treatment with auranofin induces massive changes in the redox state of the proteome. **b,** Waterfall plot of the top targets ranked across all three dimensions upon auranofin treatment (all proteins have a Fisher *p* value < 0.05 across all three dimensions). The pie piece size is proportional to the ranking of each protein in the respective dimension. **c,** Radarplots depict the redox, solubility and expression changes of top proteins GCLM and TXNRD1 upon auranofin treatment (the radarplots range from log2 fold change -2.5 to 2.5). **d,** Correlation of protein abundance changes calculated using actual expression channels or those inferred from the non-Cys peptides in REX (REXpression) in dataset 2. Only peptides passing significance in REXpression were used. **e,** The distribution of CV between replicates for REXpression vs. expression in dataset 2. **f.** Correlation between fold changes obtained from normalization of REX data to the expression channels vs. to the sum of non-Cys peptides from the same REX channels in dataset 2. **g,** The scatterplot of different dimensions highlights proteins changing in different dimensions (the cognate target TXNRD1 is shown in green circles).

In each dimension of analysis, *p* value was estimated using Student’s t-test with unequal variance based on the normalized intensities of treated vs. control samples. For control of the false discovery rate (FDR), a permutation analysis of the protein abundances between sample and control replicates was made. In total, 10 rounds of permutation were performed for each dimension and thus estimated average FDR was generally below 4% in all dimensions. This analysis was also performed for all the other datasets within the study.

We have previously introduced a robust ranking system for deconvolution of drug targets across multiple experiment types ^1^. In this approach, in each individual dimension, the proteins or peptides are ranked based on their fold change and, separately, based on *p* values (detailed in Methods). Then these ranks for all dimensions are summed together, and the proteins are sorted by the final ranking in ascending order. Applying to dataset 1 the same ranking system and a coefficients of variation (CV) cutoff of 30% for protein data in all dimensions, auranofin target thioredoxin reductase 1 (TXNRD1) ranked 2^nd^ out of the 3,252 proteins common for all dimensions, after only glutamate-cysteine ligase regulatory subunit (GCLM) (“waterfall plot” on **Fig. 2b**; full rankings in **Supplementary Data 3**). In our previous study on auranofin, TXNRD1 ranked 3^rd^ among the dramatically lower number of proteins common for all three dimensions (232 proteins) due to the low depth of conventional redox proteomics analysis. Nuclear factor NF-kappa-B p100 subunit (NFKB2) and cysteine and histidine-rich domain-containing protein 1 (CHORDC1) previously found on the first and second positions received in our current PISA- REX analysis the rankings 636 and 25, respectively ^1^. This discrepancy could be due to the much higher proteome coverage in the current study. The pie piece sizes in **Fig. 2b** show the contribution of each dimension to the ranking of a protein of interest. For example, for TXNRD1, the most decisive dimension propelling the protein to the top of the list was solubility provided by PISA analysis. Radar plots in **Fig. 2c** depict the changes of the top protein GCLM and TXNRD1 across the three dimensions.

The inhibition of TXNRD1 is known to induce the NFE2L2 (nuclear factor erythroid 2- related factor 2) or NRF2 pathway ^29^, which then activates oxidative stress response genes such as GCLM and glutamate-cysteine ligase catalytic subunit (GCLC), that are involved in glutathione (GSH) synthesis ^30^. In agreement with that and our previous results ^1,31^, the top 30 proteins mapped to NRF2 pathway (TXNRD1, GCLM, GCLC, PRDX1 and SQSTM1), glutathione synthase complex (GCLM and GCLC) and selenocysteine pathway (TXNRD1, GCLM, GCLC and PRDX1). These results provide the proof of principle for the PISA-REX approach.

To test whether protein abundances can also be deduced from the REX channels, thus allowing for a more compact experiment, we performed an additional LC-MS/MS analysis (dataset 2), where PISA, redox and expression dimensions were all profiled after 24 h of treatment at IC50 concentration of auranofin (**Supplementary data 4** and **5** corresponding to protein and peptide level data). In this dataset 2, we quantified 92,023 peptides in total. Of the 18,931 Cys containing peptides, 17,444 carbamidomethylated molecules were quantified belonging to 3,921 proteins. Expression fold changes (auranofin vs. control) calculated based on the abundance information derived from non-Cys containing peptides (hereafter called REXpression) correlated well with those calculated based on the actual expression channels (R=0.91) (**Fig. 2d**). The CVs of the conventional protein expression and REXpression were also very similar (**Fig. 2e**), both in terms of the median value (8% and 7%, respectively) as well as mean values (12% and 10%, respectively). Note that both parameters here are in favor of REXpression, which was also noted for the other datasets (see below).

In combined REXpression ranking (24 h, dataset 2) with PISA and redox rankings (2 h, dataset 1), TXNRD1 retained one of the top positions among 2,814 proteins (full rankings in **Supplementary Data 6**). Furthermore, a good correlation (R=0.89) was observed between the redox data normalized by REXpression vs. those normalized by actual expression channels (**Fig. 2f**). Therefore, we concluded that the PISA-REX experiment can be further compacted by excluding the expression channels and using REXpression instead. Since such an approach leads in general to lower CVs, throughout this paper we will use REXpression data for protein abundances wherever applicable, even when conventional expression is available in “all-Cys channels” as well.

We then investigated the correlations between the three dimensions of analysis. **Fig. 2g** shows only weak correlations among them in our second PISA-REX dataset, proving the orthogonality of these dimensions for the proteins not involved in drug action mechanism. But the outliers in these plots are not random proteins. As we have shown before, protein solubility/stability changes can be linked to oxidation and reduction of Cys residues ^1^. Thus, the outliers in the PISA vs. REX scatter plot could be rich with mechanistic proteins. Consistent with that, the cognate target of auranofin TXNRD1 was among the significantly shifting proteins in two dimensions. Furthermore, thioredoxin-like protein 1 (TXNL1) was the most significant outlier in all three dimensions. Our previous studies have shown extreme downregulation of TXNL1 upon auranofin treatment ^31^ and concluded that this protein might be a substrate of TXNRD1 ^32^. Here we found that the redox state of TXNL1 is also modulated in response to TXNRD1 inhibitor auranofin.

Consistent with the earlier observations that detection of the less abundant oxidized Cys-peptides is more challenging, the numbers of detected peptides modified with NEM was substantially lower than the number of IAA-modified peptides (1,848 vs 17,089 in dataset 1 and 2,458 vs 17,444 in dataset 2). Somewhat surprisingly, the oxidation ratios based on the NEM-modified peptides did not anti-correlate well with those calculated based on carbamidomethylated peptides, and this result was consistent for all the analyzed datasets in the paper. The lack of the statistical link could be due to the poor accuracy in the measurement of low-abundant NEM peptides as well as the presence of many different Cys modifications that can distort the estimation of oxidation ratios based on merely NEM-modified peptides.

### Interferons remodel the monocyte proteome in all three dimensions

Next we applied PISA-REX to investigate proteome remodeling by IFNs, signaling proteins that constitute the first line of defense against viral infections in mammals ^33^. IFNs interfere with viral replication by activating immune cells through induction of transcription of hundreds of IFN-stimulated genes (ISGs) leading to a massive proteome response ^34^. A recent study has applied expression proteomics and protein correlation profiling in combination with size exclusion chromatography to investigate the dynamic rewiring of the human interactome by IFN-β signaling ^35^. However, to date, no study has been performed on the IFN-induced proteome rewiring in the redox and protein solubility/stability dimensions. Therefore, here we applied PISA-REX to THP1 monocytes treated with IFN-α. As IFNs are known to induce oxidative stress ^36^, we assessed ROS production across a series of IFN-α concentrations. Upon 10 ng/mL IFN-α treatment, we observed a ∼30% increase in ROS production compared to untreated controls, without any discernible effect on cell viability (**Fig. 3a**). For PISA-REX analysis we chose this concentration, which was also used in previous studies ^35^. The PISA-REX data for the protein and peptide levels on THP1 cells treated with IFN-α for 16 h can be found in **Supplementary data 7** and **8**. 16,076 peptides were carrying the carbamidomethyl modification (12,957 unique sequences), mapping to 4,315 proteins.

**Fig. 3.**
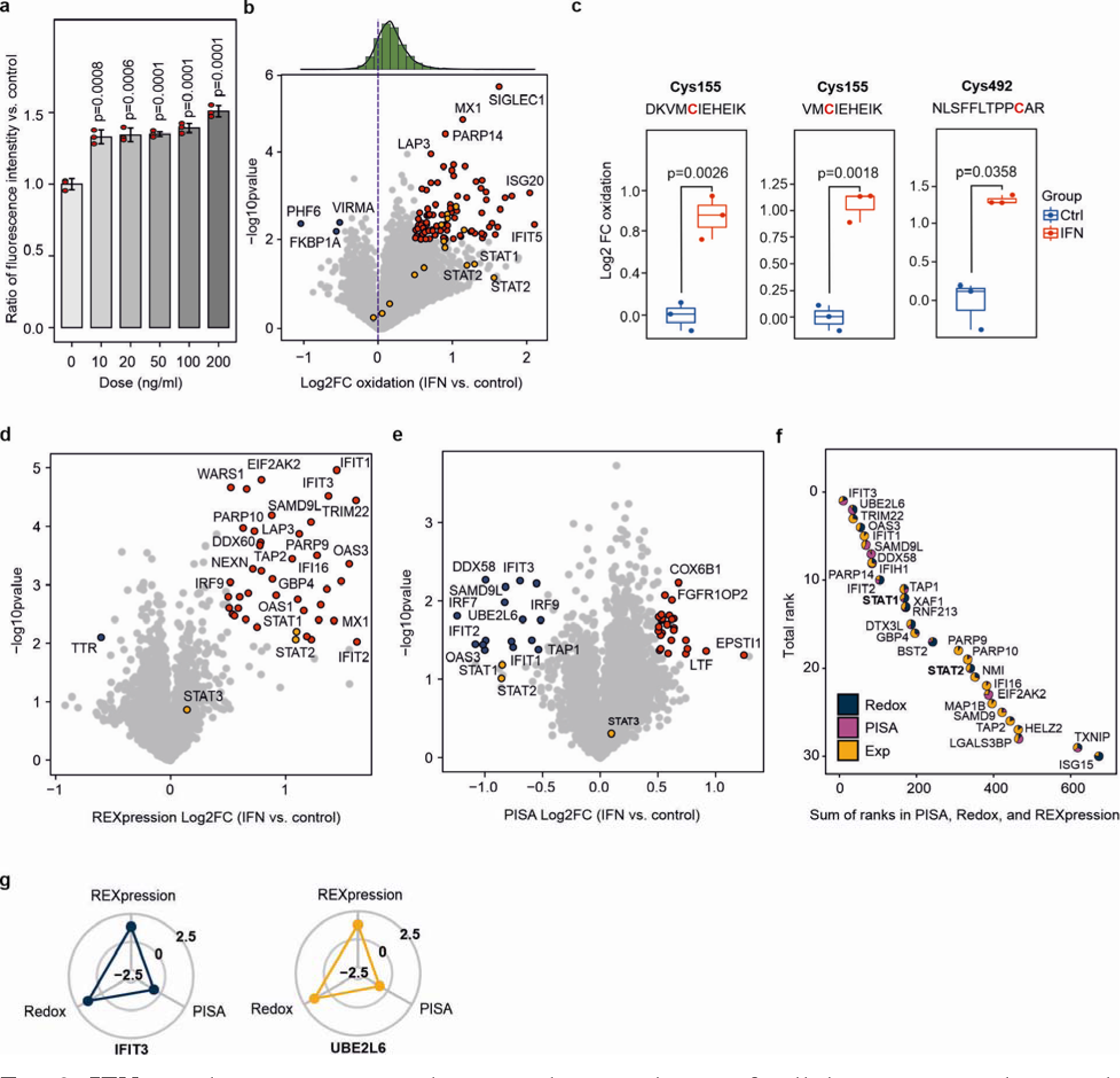
IFN-α induces massive oxidation and upregulation of cellular proteins, along with changes in protein solubility. **a,** The extent of ROS generation across different IFN-α concentrations (n=3 independent biological replicates; mean±SD; two-sided Student’s t-test). **b,** Redox changes in the proteome upon IFN-α treatment. **c,** The oxidation of three STAT1 peptides upon IFN-α treatment. Significance is calculated by two-sided Student’s t-test with unequal variance. Boxplots: Center line-median; box limits contain 50% of data; upper and lower quartiles, 75 and 25%; maximum-greatest value excluding outliers; minimum-least value excluding outliers; outliers-more than 1.5 times of the upper and lower quartiles. Expression (**d**) and solubility/solubility (**e**) changes in the proteome upon IFN-α treatment. **f,** Top ranking proteins across all three dimensions (all proteins have a combined Fisher *p* value < 0.05). The pie piece size is proportional to the ranking of each protein in the respective dimension – larger sector for higher ranking. **g,** The change in the redox state, solubility, and expression of top-ranking proteins. The radarplots range from log2 fold change -2.5 to 2.5.

As expected, IFN-α treatment induced massive oxidation of cellular proteins, denoted by the asymmetry of the density plot in **Fig. 3b**. Some of the top proteins, such as STAT1, XAF1, RNF123 and PRDX4 were already known to be redox-related. For instance, the redox state of Cys324 and Cys492 residues in STAT1 can be regulated by S-glutathionylation ^37^. Here we confirmed the oxidation of Cys492 and additionally found oxidation of Cys155 (**Fig. 3c**), supported by observation of five peptides (with two different sequences) containing this Cys. Encouragingly, our redox proteomics approach has estimated very similar oxidation ratios of the two peptides with different sequences covering Cys155 (**Fig. 3c**). In XAF1 that is known to be transcriptionally activated by oxidative stress ^38^, we found oxidation of Cys112 upon IFN stimulation. As expected, the IFN-α treatment led to changes in the REXpression and solubility of many known and putative ISGs (**Fig. 3d-e**; the scatterplots of the three dimensions are shown in **Supplementary Fig. 1a)**. The top 30 proteins ranked across the three dimensions are shown in **Fig. 3f** (full rankings in **Supplementary Data 9**). The top protein out of 3,757 proteins shared among the three dimensions was IFIT3, interferon-induced protein with tetratricopeptide repeats 3. The 5^th^ top protein was IFIT1, interferon induced protein with tetratricopeptide repeats, while the 8^th^ was IFIH1, interferon induced protein with helicase C domain 1. STAT1 and STAT2, transcription factors of IFN response ^39,40^, ranked 12^th^ and 20^th^, respectively. At least 14 of the top 30 proteins mapped to IFN signaling (p=4.62e-17). For these two top proteins, redox and REXpression were the two defining dimensions (**Fig. 3g**).

Similar to the auranofin dataset, REXpression fold changes here correlated well (R=0.94) with the fold changes obtained from the actual expression channels (**Supplementary** Fig. 1b). The CVs between the replicates for REXpression were again lower than for expression (10.3% vs 12.3% for medians, and 12.8% vs 15.8% for average values), confirming that REXpression gives a more precise estimate for protein abundance changes.

### pDCs with *Ncf1*-mutation respond differently to IFN-**α**

Next, we applied PISA-REX to study the role of NCF1 in a murine model of systemic lupus erythematosus (SLE). SLE is a chronic autoimmune disease predominantly observed in women of child-bearing age, affecting 0.1% of global population ^41^. SLE patients manifest overactivated type I IFN signature and high expression of ISGs, which are reflective of disease activity and severity ^41^. ROS is considered to have a regulatory role in the development of SLE, similar to other inflammatory and autoimmune diseases ^42^.

Neutrophil cytosol factor 1 (NCF1) is a subunit of the NADPH oxidase 2 (NOX2) complex, which plays a crucial role in ROS production to protect against invading pathogens. The dysfunctional *NCF1*-339 allele, which impairs the function of NOX2 complex, is the major genetic association for SLE, with regards to odds ratio and allelic frequency ^43,44^. Dysfunctional NCF1 affects ROS production and is strongly associated with SLE in humans and mouse models ^45^. Our previous study has found plasmacytoid dendritic cells (pDCs) as the key immune cell type driving SLE pathogenesis ^45^. ROS deficiency enhances the generation, accumulation and function of pDCs, which exacerbates pristane-induced and spontaneous lupus. IFN-α is known to autocrinally control pDC functions in cellular development, differentiation, maturation and survival ^46,47^.

To uncover the molecular phenomena by which ROS affects pDC function in NCF1- dependent development of lupus, we isolated pDCs from the bone marrow-derived cells of wild type (wt) vs. *Ncf1*-mutant (*Ncf1^m1j/m1j^*) B10.Q mice. Subsequently, the cells were stimulated with 500U/mL IFN-α for 20 h in culture. Since the cells were derived using magnetic bead sorting with anti-B220 antibody, the sorted population comprised mature pDCs and CCR9^-^pDC precursors, both exhibiting strong IFN-α responses ^48^. Due to the point mutation in the intron7 of *Ncf1* in our model, p47phox_Δ228–235 is expressed at a low level with 8 amino acid residues being deleted, leading to functional impairment of NCF1 function ^49^.

The PISA-REX data corresponding to protein and peptide levels can be found in **Supplementary Data 10** and **11**. In total, 9670 carbamidomethylated peptides were quantified without missing values. The scatterplots of different dimensions are shown in **Fig. 4a-c**. The overall oxidation rate was higher in the *Ncf1*-mutant cells vs wt, indicating that the mutation affects response to IFN-α treatment on the redox level (**Fig. 4a**). The median CVs for REXpression and expression were 8.1% and 8.6%, respectively (corresponding averages being 10.3% and 10.5%). The waterfall plot of the top-ranking proteins is shown in **Fig. 4d** (full rankings in **Supplementary Data 12**). In the absence of 8 residues in the NCF1 SH3_B_ domain, which impairs NCF1 translocation to the membrane, significant changes in the solubility, expression and the redox state of this protein were induced under IFN-α treatment conditions (**Fig. 4e**). Interestingly, NCF2, another member of the NOX2 complex, behaved similarly. Three different NCF1 Cys residues were reduced in *Ncf1*-mutant pDC cells upon IFN stimulation compared to wt cells (**Fig. 4f**).

**Fig. 4.**
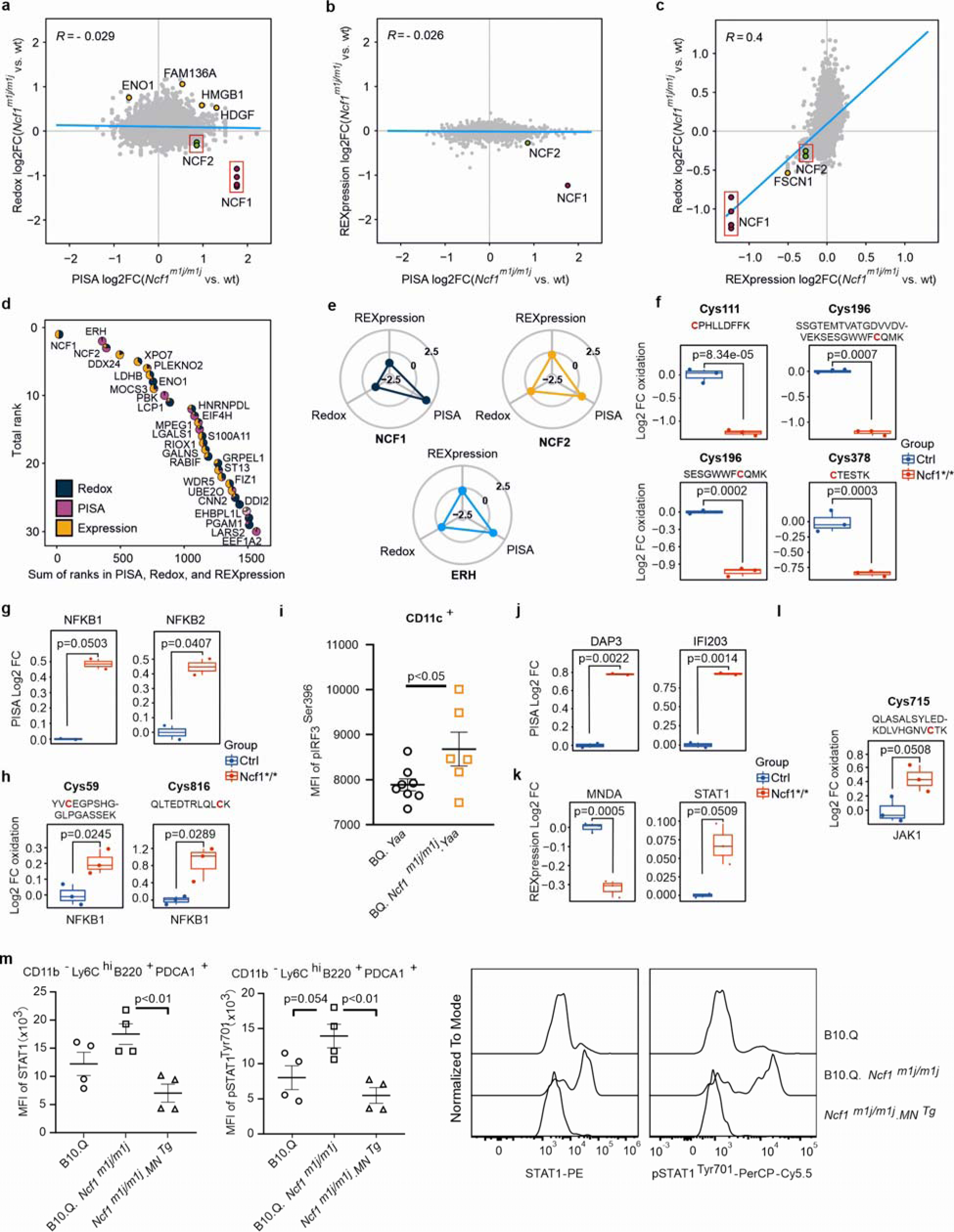
PISA-REX for studying a mouse model of lupus. **a-c,** Changes in the redox state, solubility and expression level of proteins in PDC cells isolated from *Ncf1^m1j/m1j^* mice vs. wt littermates stimulated with IFN-α in cell culture. **d,** The top targets ranked across three dimensions. The pie piece size is proportional to the ranking of each protein in the respective dimension. **e,** The changes in the redox state, solubility and expression level of top targets. The radarplots range from log2 fold change -2.5 to 2.5. The redox state of four NCF1 peptides (**f**), the change in the solubility of NFKB1 and NFKB2 (**g**), and redox state of NFKB1 (**h**) in *Ncf1*- mutant vs. wt cells stimulated with IFN. **i,** Phosphorylation of IRF3 in dendritic cells within peripheral blood from 3-month-old BQ.*Yaa* (n=8) and BQ.*Ncf1^m1j/m1j^.Yaa* (n=6) mice. The differential solubility (**j**), REXpression (**k**) and redox state (**l**) of proteins as representative ISGs, in *Ncf1*-mutant vs. wt cells stimulated with IFN. **m,** Expression of STAT1 and p-STAT1 in pDCs within peritoneal exudate cells from B10.Q (n=), B10.Q.*Ncf1^m1j/m1j^*, and *Ncf1^m1j/m1j^*.MN^tg^ mice (of each n=4) at day 3 post pristane injection. Representative histograms are presented. Results are shown as mean ± SEM. Statistical significance is determined by Two-tailed Mann-Whitney U test in (**i**) and one-way analysis of variance with Tukey’s multiple comparison test in (**m**). For all other panels, significance is calculated by two-sided Student’s t-test with unequal variance. Boxplots: Center line-median; box limits contain 50% of data; upper and lower quartiles, 75 and 25%; maximum-greatest value excluding outliers; minimum-least value excluding outliers; outliers-more than 1.5 times of the upper and lower quartiles

Earlier studies have shown that pDCs with the *Ncf1* mutation produce higher levels of IFN-α, partially through the STING pathway ^45^. Upon stimulation by cytosolic DNA, STING recruits NF-kB and IRF3 to undergo phosphorylation and activation by TBK1, leading to the production of type I IFNs and other cytokines ^50^. PISA-REX data analysis showed that upon IFN stimulation, the solubility of NFKB2 and NFKB1 was higher in *Ncf1*-mutant cells vs wt (**Fig. 4g**) and Cys59 and Cys816 from NFKB1 were more oxidized (**Fig. 4h**). While the loss of IRF3 solubility in PISA was not statistically significant, we observed increased phosphorylation of IRF3 in dendritic cells from peripheral blood of *Ncf1*-mutant mice in the spontaneous lupus model (**Fig. 4i**).

The *Ncf1* mutation was also found to regulate the downstream signals mediated by IFN- α. For example, *Ncf1* mutation status affected the solubility of DAP3 and IFI203 (**Fig. 4j**), expression of MNDA and STAT1 (**Fig. 4k**), and redox state of NFKB1 (**Fig. 4h**) as well as JAK1 (**Fig. 4l**). Our data confirm the regulation of STAT1, a transcriptional factor for ISGs, at the early stage of pristane-induced lupus. PDCs from *Ncf1*-mutant mice exhibit higher levels of STAT1 expression and phosphorylation compared to wt mice, while the levels in *Ncf1^m1j/m1j^*.MN^Tg^ mice, where Ncf1 is specifically restored in pDCs and monocytes/macrophages, are comparable to wt (**Fig. 4m**).

These results are consistent with our previous finding that NCF1 deficiency enhances the JAK1/STAT1 pathway, resulting in higher expression of ISGs and promoting lupus in both mice and humans ^43,45^. A curios observation is the behavior of the “enigmatic” protein Enhancer of Rudimentary Homolog or ERH ^51^ that largely mimics that of NCF1 and NCF2 (**Fig. 4e**). Erh is involved in protein complexes related to pyrimidine metabolism, acts as a transcriptional repressor, and participates in cell cycle regulation ^52^. The involvement of this protein in autoimmune diseases warrants further studies.

## Discussion

Here, we developed a novel redox proteomics strategy that provides 6-9 times higher multiplexing, ∼25x lower cost per sample (compared to iodoTMT labeling) and an unprecedented coverage of the cysteinome. Importantly, redox analysis using REX had an average overall CV below 14%, which is comparable to other types of proteomics analyses ^53,54^. It should also be considered that variation in the measurement of peptides is generally higher than proteins, where abundance information is obtained from multiple peptides. Furthermore, we showed that the non-Cys containing peptides in the REX channels can be used to extract protein abundance information (REXpression) that highly correlates with routine expression analysis. Furthermore, fold changes calculated based on REXpression gave consistently lower CVs than those calculated from expression, which might be explained by the complex modification landscape of Cys residues that could potentially skew protein abundances in the expression channels. REXpression will be particularly useful in future applications such as redox proteomics on small sample amounts, and can potentially be used in single cell proteomics ^55,56^.

Since REX can provide simultaneous redox and expression level information, only 12 TMT channels can accommodate the PISA-REX analysis in 3 replicates. For even more robust statistics, PISA and REX can each be performed in 4 replicates for 2 conditions using TMTpro- 16plex.

Upon benchmarking REX against our comprehensive chemical proteomics on auranofin ^1^, we integrated this tool with PISA, obtaining a possible industry standard proteomics tool. We applied such a PISA-REX tool to several model systems, showcasing target deconvolution for drugs and biological agents such as IFNs and disease mechanism in a murine model of lupus. We showed 3D proteome remodeling upon IFN-α treatment, revealing ISGs that are redox modulated and those that undergo solubility transitions on top of expression changes. By applying REX to a murine model of lupus, we demonstrated that upon IFN-α stimulation, mutant *Ncf1*, an important gene with polymorphisms in the etiology of this disease, is strongly modulated at the redox, solubility and expression levels compared to the wt mice. Furthermore, we showed the involvement of NFKB and STAT1 signaling in the molecular events in pathways dependent of NCF1. ROS deficiency enhances the generation, accumulation and function of pDCs, which are the key immune cell type driving SLE pathogenesis. Here, we uncovered the molecular pathways by which IFN-α controls pDC functions.

In this paper, we only focused on the quantification of the dominant reduced form of Cys. As the forms of Cys oxidation upon different treatments can vary, enrichment strategies might be needed for analysis of each specific type of Cys oxidation, as previously described for nitrosylation ^57^. We envision the widespread use of PISA-REX in academia and industry as a robust tool for target deconvolution as well as gaining biological insight into disease mechanisms.

## Online methods

### Cell culture

Human colorectal carcinoma HCT116 (ATCC, USA) cells were grown at 37 °C in 5% CO_2_ using McCoy’s 5A modified medium (Sigma-Aldrich, USA) supplemented with 10% FBS superior (Biochrom, Berlin, Germany), 2 mM L-glutamine (Lonza, Wakersville, MD, USA) and 100 units/mL penicillin/streptomycin (Gibco, Invitrogen). Human THP1 cells (ATCC, USA) were grown under the exact same conditions in RPMI. Low-number passages were used for the experiments and cells were checked for mycoplasma contamination.

### Viability assay

The viability assays were performed using CellTiter-Blue (Promega) according to our published protocol ^31^. Briefly, cells were seeded at a density of 4,000 per well in 96 well plates. Adherent cells were grown for one day and treated with serial concentrations of the compounds on the next day, while the suspension-type cells were treated 2 h after seeding. IC50 was determined as the concentration inducing 50% reduction in viability.

### ROS production assay

The assay was performed according to our recent protocol ^58^. Briefly, THP1 cells were cultured at a density of 250,000 cells in 6-well plates and stimulated separately in triplicates with 0, 10, 20, 50, 100 and 200 ng/mL IFNα for 16 h. After the treatment, cells were washed twice with PBS and resuspended in 2 mL PBS, and treated with 2′,7′-dichlorofluorescin diacetate (DCF-DA) to a final concentration of 20 μM and kept at 37 °C for 30 min. DCF-DA was then discarded, cells were washed twice with PBS and resuspended in 1 mL PBS again. An equal volume of cell suspension from each sample was transferred to flat bottom black 96-well plates and the fluorescence was recorded at an excitation of 495 nm and emission of 527 nm. Trypan blue dye exclusion assay was also used to count the number of living cells for normalization.

### PISA-REX experiments

Cells were treated in 8 replicates with the vehicle or compounds according to the indicated concentrations and durations in **Supplementary Table 1** for different experiments. For all-Cys and reduced Cys analyses, cells were lysed with 1% SDS, 8 M urea, Tris buffer pH 8.0 plus protease inhibitor. The cell lysates were subjected to 1 minLJsonication using Branson probe sonicator with 3LJs on/off pulses and a 30% amplitude. Protein concentration was measured using Pierce BCA Protein Assay Kit (Thermo), and the volume corresponding to 25 µg of protein was transferred from each sample to new tubes. For all-Cys analysis, DTT was added to a final concentration of 10 mM and samples were incubated for 45 min at room temperature (RT).

Subsequently, IAA was added to a final concentration of 50 mM and samples were incubated at RT for 1 h in the dark. The reaction was quenched by adding an additional 10 mM of DTT. Proteins were precipitated by methanol/chloroform and resuspended in 20 mM EPPS buffer pH 8.5 with 8M urea.

For reduced Cys samples, IAA was added to a final concentration of 50 mM and samples were kept in the dark for 1 h at RT. The proteins were then precipitated by chloroform and resuspended in the same volume of lysis buffer again. DTT was added to the final concentration of 10 mM and samples were incubated for 30 min at RT. NEM was added at the final concentration of 50 mM, followed by 1 h incubation in the dark at RT. After a further precipitation step, the samples were resuspended in 20 mM EPPS buffer pH 8.5 with 8M urea.

For PISA in intact cells, we followed our previously published protocol ^4,26^. Cells were collected, centrifuged, and washed twice with PBS and then resuspended in ∼300 μL PBS. The cell suspension from each replicate were aliquoted into 10 in 96-well plates and heated in an Eppendorf gradient thermocycler (Mastercycler X50) in the temperature range of 48-59 °C for 3 min. Samples were cooled for 3 min at RT and afterwards snap frozen with liquid nitrogen and kept on ice. The cells were then lysed by 5 times freezing in liquid nitrogen and thawing at RT. Samples from each replicate were then combined and transferred into polycarbonate thickwall tubes and centrifuged at 100,000 g and 4 °C for 20 min. The soluble protein fraction was then collected and subjected to protein concentration measurement using the BCA assay. The reduction and alkylation were performed as detailed for all-Cys samples. After precipitation, PISA samples were also resuspended in 20 mM EPPS buffer pH 8.5 with 8M urea.

The rest of the protocol is the same for all three types of samples and adapted from our previous protocol ^31^. Urea was diluted to 4M by adding 20 mM EPPS. Lysyl endopeptidase (LysC; Wako) was added at a 1:75 w/w ratio and incubated at RT overnight. Samples were diluted with 20 mM EPPS to the final urea concentration of 1M, and trypsin was added at a 1:75 w/w ratio, followed by incubation for 6 h at RT.

Acetonitrile (ACN) was added to a final concentration of 20% and TMT reagents were added 4x by weight (200 μg) to each sample, followed by incubation for 2 h at RT (TMT channel assignment information is in **Supplementary Table 2**). The reaction was quenched by addition of 0.5% hydroxylamine. Samples within each replicate were combined, acidified by TFA, cleaned using Sep-Pak cartridges (Waters) and dried using DNA 120 SpeedVac Concentrator

(Thermo). The pooled samples were resuspended in 20 mM ammonium hydroxide and separated into 96 fractions on an XBrigde BEH C18 2.1x150 mm column (Waters; Cat#186003023), using a Dionex Ultimate 3000 2DLC system (Thermo Scientific) over a 48 min gradient from 1% to 63% B (B=20 mM ammonium hydroxide in acetonitrile) in three steps (1-23.5% B in 42 min, 23.5-54% B in 4 min and then 54-63%B in 2 min) at 200 µL min^-1^ flow. Fractions were then concatenated into 24 samples in sequential order (*e.g.* A1, C1, E1 and G1 on the 96 well plate were combined).

### LC-MS/MS

After drying, samples were dissolved in buffer A (0.1% formic acid and 2% ACN in water). The samples were loaded onto a 50 cm EASY-Spray column (75 µm internal diameter, packed with PepMap C18, 2 µm beads, 100 Å pore size) connected to a nanoflow Dionex UltiMate 3000 UHPLC system (Thermo) and eluted in an organic solvent gradient increasing from 3-4% to 26-28% (B: 98% ACN, 0.1% FA, 2% H_2_O) at a flow rate of 300 nL min^-1^ over 95 min. The eluent was ionized by electrospray, with molecular ions entering an Orbitrap Fusion Lumos mass spectrometer (Thermo Fisher Scientific) for auranofin samples. The rest of the samples were analyzed on either Orbitrap HF, Exploris or Lumos (all - Thermo Fisher Scientific). Method settings are tabulated in **Supplementary Table 3**.

### Data processing

The raw LC-MS/MS data were analyzed by Proteome Discoverer v2.5 (Thermo Fisher Scientific) using Mascot Server v2.5.1 as search engine with SwissProt human (20,360 entries downloaded on 28 February 2022) or mouse (21,989 entries downloaded on 2 September 2020) and common contaminant databases. No more than two missed cleavages were allowed. A 1% false discovery rate was used as a filter at both protein and peptide levels. For all other parameters, the default settings were used. In all analyses, reporter ions of TMTpro 16-plex were used for peptide quantification. Cys carbamidomethylation and N-ethylmaleimide, methionine oxidation, asparagine and glutamine deamidation, N-terminal acetylation and TMTpro (+304.207 Da) were selected as dynamic modifications. Contaminants were removed. A mutation in the Ncf1 gene (m1j) leads to a truncated NCF1 protein, thereby a non-functional NOX2 complex, as described before ^49^. In the *Ncf1* dataset, no peptides containing the mutation site could be quantified; therefore, for unbiased comparison of the mutant with the wt samples, we removed the two peptides mapping to the sequence containing this site and calculated the protein abundances based on the other 12 detected peptides.

### Animals

C57BL/6NJ mice were originally from The Jackson Laboratory (Farmington, CN, USA). B6N.Q (C57/B6N.Q/rhd) and B10.Q (C57/B10N.Q/rhd) have been fully backcrossed into B6N and B10 genomes but with the MHC region from DBA/1. A mutation in the *Ncf1* gene (*m1j*) leads to a truncated NCF1 protein, thereby a non-functional NOX2 complex, as described before ^49^. The derived *Ncf1*-mutant mouse strains on different backgrounds were designated as B6N.Q.*Ncf1^m1j/m1j^* and B10.Q.*Ncf1^m1j/m1j^*, respectively. The transgenic strain B10.Q.*Ncf1^m1j/m1j^.MN^Tg^*(herein denoted as *Ncf1^m1j/m1j^.MN^Tg^*) express functional NCF1 restricted to macrophages/monocytes using the human CD68 promoter ^59^. B6.SB-*Yaa*/J (Stock No: 000483) from The Jackson Laboratory has been fully backcrossed onto B10.Q, denoted as B10.Q.*Yaa*. The *Yaa*-carrying strain with functional NCF1 (BQ.*Yaa*) or mutant NCF1 (BQ.*Ncf1^m1j/m1j^*.*Yaa*) was obtained by crossing B6N.Q.*Ncf1^m1j/m1j^* with B10.Q.*Yaa*. Mice were housed under specific pathogen-free (FELASAII) conditions in individual ventilated cages with wood shaving bedding in a climate-controlled environment with a 12 h light/dark cycle. Animal experiments were performed in a controlled way balanced for age and sex; the genotype of the mice was blinded to the investigators, following the ARRIVE guidelines.

### PDC isolation and stimulation

BM cells were flushed from 2 femurs and 2 tibias per mouse. After lysis of red blood cells with Ammonium-Chloride-Potassium buffer for 1 min at RT, cells were plated at 2×10^6^ cells/mL in RPMI complete media containing 200ng/mL FLT3L (BioLegend, #550704) for pDC differentiation. Cells were collected on day 9 and pDCs were magnetically isolated with biotinylated anti-B220 mAb (BD Biosciences, clone: RA3-6B2). Sorted pDCs were cultured at a density of 5×10^6^ cells/well in RPMI medium and stimulated with 500U/mL IFN-α (BioLegend, #752804) for 20 h.

### Pristane-induced lupus

B10.Q, B10.Q.*Ncf1^m1j/m1j^*, and *Ncf1^m1j/m1j^*.MN^Tg^ mice were injected intraperitoneally with a single dose of 500 μL pristane (Sigma-Aldrich, #P2870).

### Flow cytometry

Single-cell suspensions were prepared from peritoneal exudates and peripheral blood from mice. After a homemade anti-mouse CD16/CD32 FcR block (clone: 2.4G2), the cells were incubated with LIVE/DEAD™ Fixable Near-IR Dead Cell Stain Kit (ThermoFisher, #L10119) and labeled with the following antibodies: anti-CD11c (BD Biosciences, clone: HL3, PE- Cyanine7), anti-CD11b (BioLegend, clone: M1/70, Pacific Blue), anti-Ly6C (BioLegend, clone: HK1.4, FITC), anti-PDCA1 (BioLegend, clone: 927, Alexa Fluor 647), anti-B220 (BD Biosciences, clone: RA3-6B2, PE-Cyanine7), anti-STAT1 (BD Biosciences, clone: 1/Stat1, PE), anti-pSTAT1^Tyr701^ (BD Biosciences, clone: 4a, PerCP/Cy5.5), and anti-pIRF3^Ser396^ (Bioss Antibodies, polyclonal, FITC). For intranuclear staining, cells were fixed and permeabilized with eBioscience Foxp3 / Transcription Factor Fixation/Permeabilization Kit (ThermoFisher, #00- 5521-00). Samples were acquired using ThermoFisher Attune NxT Flow Cytometer equipped with ThermoFisher Attune NxT Software version 3.2.1, and the data were analyzed using the BD FlowJo™ version 10.7.1.

### Statistical Analysis

Data analysis was performed using R project versions 3.6 - 4.0. The raw abundances were normalized to the total channel intensity. Protein abundances for PISA were normalized to protein abundances when the treatment times were matched. Redox data was analyzed on the peptide level, where the abundance of a given peptide was calculated by dividing its intensity to the summed intensity of that channel for a given sample. For redox analysis thus obtained abundance of Cys-containing peptide was normalized by the summed abundance of non-Cys peptides from the same protein. Oxidation ratio was calculated as the normalized abundance of peptide in control vs. treated samples (in NEM-modified peptide analysis, calculation of oxidation ratio was reverse). *P* values for the potential target proteins were calculated by t-test based on normalized abundances between control and treatments in all cases. Two-sided t-test with unequal variance was applied to calculate *p* values, unless otherwise specified. For calculation of statistical significance across the three dimensions, Fisher formula was used for calculation of the merged *p* values.

For target ranking, the proteins were individually ranked in each dimension for absolute values of the log2 of their abundance ratio (fold change, FC) and the *p* value and then the sum of the two ranks was calculated. For expression, ranking was done on the log2 FC, as the majority of targets are up-regulated ^1,31^. Then thus obtained rankings were summed across the three dimensions, and the proteins were ranked again.

### Network mapping

GO pathway enrichment analysis for the top proteins was performed using the StringDB tool version 12 ^60^.

### Data availability

The LC-MS/MS raw data files and extracted peptides and protein abundances are deposited in the jPOST repository of the ProteomeXchange Consortium ^61^ under the dataset identifiers PXD035964 (animal experiment) [https://www.ebi.ac.uk/pride/archive/projects/PXD035964] and PXD042967 for all the other datasets with no restrictions. The extracted protein abundances data and relevant outputs of data analysis are provided in Supplementary Data 1-12. All data are also available from the corresponding authors on reasonable request. The source data underlying **Figs. 3a**, **4i** and **4m** are provided as a Source Data file. The remaining panels are produced from the Supplementary Data tables.

## Acknowledgements

This work was supported by a grant from Cancerfonden (22 1967 Pj to R.A.Z). A.A.S. was supported by Swedish Research Council (grant 2020-00687) and Swedish Society of Medicine (grant SLS-961262, 1086 Stiftelsen Albert Nilssons forskningsfond) and KI foundation research grant (2022-02035). R.H. was supported by the Cancer foundation (grant 22 2350 Pj 01 H), Science Research Council (grant 2019-01209) and the KA Wallenberg foundation (grant 2019.0059). Jaakko S Teppo was supported by Academy of Finland (project #321472). Open access funding was provided by Karolinska Institutet.

## Author contributions

Conceptualization, R.A.Z. and A.A.S.; methodology and experiment design, A.A.S., R.A.Z., S.P.G., X.Z., H. Luo and A.V.; project organization, training, resources and funding acquisition, R.A.Z., A.A.S. and R.H.; Proteomics experiments, A.A.S., H.Lyu., J.T., and M.G.; *in vivo* experiments, H.Luo.; data analysis and visualization, A.L., A.A.S., A.V. and H.G.; writing—original draft, A.A.S. and R.A.Z.; writing—review & editing, all other co-authors.

## Supplementary Materials

Supplementary Information text file

**Supplementary Data 1.** PISA assay and expression data on HCT116 cells treated with auranofin for 2h and 48h, respectively.

**Supplementary Data 2.** Redox proteomics data on HCT116 cells treated with auranofin for 2h **Supplementary Data 3.** The ranking of targets across the three dimensions of redox state (2h), solubility (2h) and expression (48h) for HCT116 cells treated with auranofin

**Supplementary Data 4.** PISA assay, expression and REXpression data on HCT116 cells treated with auranofin for 24h

**Supplementary Data 5.** Redox proteomics data on HCT116 cells treated with auranofin for 24h **Supplementary Data 6.** The ranking of targets across the three dimensions of redox state (2h), solubility (2h) and REXpression (24h) in HCT116 cells treated with auranofin

**Supplementary Data 7.** PISA assay, REXpression and expression data on THP1 cells treated with interferon for 16h

**Supplementary Data 8.** Redox proteomics data on THP1 cells treated with interferon for 16h **Supplementary Data 9.** The ranking of targets across the three dimensions of redox state, solubility and REXpression in THP1 cells treated with interferon

**Supplementary Data 10.** PISA assay, REXpression and expression data on PDC cells derived from WT and Ncf1 mutant mice and treated with interferon

**Supplementary Data 11.** Redox proteomics data on PDC cells derived from WT and Ncf1 mutant mice and treated with interferon

**Supplementary Data 12.** The ranking of targets across the three dimensions of redox state, solubility and REXpression for PDC cells derived from Ncf1 mutant vs. WT mice and treated with interferon

## References

1. Saei, A. A. et al. Comprehensive chemical proteomics for target deconvolution of the redox active drug auranofin. Redox Biol. 101491 (2020).

2. Savitski, M. M. et al. Tracking cancer drugs in living cells by thermal profiling of the proteome. Science (80). (2014) doi:10.1126/science.1255784.

3. Molina, D. M. et al. Monitoring drug target engagement in cells and tissues using the cellular thermal shift assay. Science (80). (2013) doi:10.1126/science.1233606.

4. Gaetani, M. et al. Proteome Integral Solubility Alteration: A High-Throughput Proteomics Assay for Target Deconvolution. J. Proteome Res. (2019) doi:10.1021/acs.jproteome.9b00500.

5. Sabatier, P. et al. An integrative proteomics method identifies a regulator of translation during stem cell maintenance and differentiation. Nat. Commun. (2021) doi:10.1038/s41467-021-26879-4.

6. Li, J. et al. TMTpro reagents: a set of isobaric labeling mass tags enables simultaneous proteome-wide measurements across 16 samples. Nat. Methods (2020) doi:10.1038/s41592-020-0781-4.

7. Li, J. et al. TMTpro-18plex: The Expanded and Complete Set of TMTpro Reagents for Sample Multiplexing. J. Proteome Res. (2021) doi:10.1021/acs.jproteome.1c00168.

8. Fratelli, M. et al. Identification by redox proteomics of glutathionylated proteins in oxidatively stressed human T lymphocytes. Proc. Natl. Acad. Sci. U. S. A. (2002) doi:10.1073/pnas.052592699.

9. Laragione, T. et al. Redox regulation of surface protein thiols: Identification of integrin α- 4 as a molecular target by using redox proteomics. Proc. Natl. Acad. Sci. U. S. A. (2003) doi:10.1073/pnas.2434516100.

10. Wojdyla, K. & Rogowska-Wrzesinska, A. Differential alkylation-based redox proteomics - Lessons learnt. *Redox Biology* at 10.1016/j.redox.2015.08.005 (2015).

11. Kim, H. J., Ha, S., Lee, H. Y. & Lee, K. J. Rosics: Chemistry and proteomics of cysteine modifications in redox biology. Mass Spectrom. Rev. (2015) doi:10.1002/mas.21430.

12. Finkel, T. Signal transduction by reactive oxygen species. Journal of Cell Biology at 10.1083/jcb.201102095 (2011).

13. Rhee, S. G. H2O2, a necessary evil for cell signaling. Science at 10.1126/science.1130481 (2006).

14. Nietzel, T., Mostertz, J., Hochgräfe, F. & Schwarzländer, M. Redox regulation of mitochondrial proteins and proteomes by cysteine thiol switches. Mitochondrion (2017) doi:10.1016/j.mito.2016.07.010.

15. Pan, Y. et al. Quantitative proteomics reveals the kinetics of trypsin-catalyzed protein digestion. Anal. Bioanal. Chem. 406, 6247–6256 (2014).

16. Bachi, A., Dalle-Donne, I. & Scaloni, A. Redox proteomics: Chemical principles, methodological approaches and biological/biomedical promises. Chemical Reviews at 10.1021/cr300073p (2013).

17. Xiao, H. et al. A Quantitative Tissue-Specific Landscape of Protein Redox Regulation during Aging. Cell (2020) doi:10.1016/j.cell.2020.02.012.

18. Doron, S. et al. SPEAR: A proteomics approach for simultaneous protein expression and redox analysis. Free Radic. Biol. Med. (2021) doi:10.1016/j.freeradbiomed.2021.10.001.

19. García-Santamarina, S. et al. Monitoring in vivo reversible cysteine oxidation in proteins using ICAT and mass spectrometry. Nat. Protoc. (2014) doi:10.1038/nprot.2014.065.

20. Leichert, L. I. et al. Quantifying changes in the thiol redox proteome upon oxidative stress in vivo. Proc. Natl. Acad. Sci. U. S. A. (2008) doi:10.1073/pnas.0707723105.

21. Araki, K. et al. Redox sensitivities of global cellular cysteine residues under reductive and oxidative stress. J. Proteome Res. (2016) doi:10.1021/acs.jproteome.6b00087.

22. Shakir, S., Vinh, J. & Chiappetta, G. Quantitative analysis of the cysteine redoxome by iodoacetyl tandem mass tags. Anal. Bioanal. Chem. (2017) doi:10.1007/s00216-017-0326-6.

23. Weerapana, E. et al. Quantitative reactivity profiling predicts functional cysteines in proteomes. Nature (2010) doi:10.1038/nature09472.

24. Yang, J. et al. Global, in situ, site-specific analysis of protein S-sulfenylation. Nat. Protoc. (2015) doi:10.1038/nprot.2015.062.

25. Van Der Reest, J., Lilla, S., Zheng, L., Zanivan, S. & Gottlieb, E. Proteome-wide analysis of cysteine oxidation reveals metabolic sensitivity to redox stress. Nat. Commun. (2018) doi:10.1038/s41467-018-04003-3.

26. Saei, A. A., et al. Mapping the GALNT1 substrate landscape with versatile proteomics tools. *bioRxiv* (2022).

27. Zhang, X. et al. Repurposing of auranofin: Thioredoxin reductase remains a primary target of the drug. Biochimie (2019) doi:10.1016/j.biochi.2019.03.015.

28. Chernobrovkin, A., Marin-Vicente, C., Visa, N. & Zubarev, R. A. Functional Identification of Target by Expression Proteomics (FITExP) reveals protein targets and highlights mechanisms of action of small molecule drugs. Sci. Rep. (2015) doi:10.1038/srep11176.

29. Cebula, M., Schmidt, E. E. & Arnér, E. S. J. TrxR1 as a potent regulator of the Nrf2- Keap1 response system. Antioxidants and Redox Signaling at 10.1089/ars.2015.6378 (2015).

30. Malhotra, D. et al. Global mapping of binding sites for Nrf2 identifies novel targets in cell survival response through chip-seq profiling and network analysis. Nucleic Acids Res. (2010) doi:10.1093/nar/gkq212.

31. Saei, A. A. et al. ProTargetMiner as a proteome signature library of anticancer molecules for functional discovery. Nat. Commun. (2019) doi:10.1038/s41467-019-13582-8.

32. Saei, A. A. et al. System-wide identification and prioritization of enzyme substrates by thermal analysis. Nat. Commun. (2021).

33. Katze, M. G., He, Y. & Gale, M. Viruses and interferon: A fight for supremacy. Nature Reviews Immunology at 10.1038/nri888 (2002).

34. Schneider, W. M., Chevillotte, M. D. & Rice, C. M. Interferon-stimulated genes: A complex web of host defenses. Annual Review of Immunology at 10.1146/annurev-immunol-032713-120231 (2014).

35. Kerr, C. H. et al. Dynamic rewiring of the human interactome by interferon signaling. Genome Biol. (2020) doi:10.1186/s13059-020-02050-y.

36. Seifert, U. et al. Immunoproteasomes preserve protein homeostasis upon interferon-induced oxidative stress. Cell (2010) doi:10.1016/j.cell.2010.07.036.

37. Butturini, E. et al. S-glutathionylation exerts opposing roles in the regulation of STAT1 and STAT3 signaling in reactive microglia. Free Radic. Biol. Med. 117, 191–201 (2018).

38. Lee, K.-W. et al. XAF1 drives apoptotic switch of endoplasmic reticulum stress response through destabilization of GRP78 and CHIP. Cell Death Dis. 13, 655 (2022).

39. Heim, M. H., Kerr, I. M., Stark, G. R. & Darnell, J. E. Contribution of STAT SH2 groups to specific interferon signaling by the Jak-STAT pathway. Science (80-.). (1995) doi:10.1126/science.7871432.

40. Wesoly, J., Szweykowska-Kulinska, Z. & Bluyssen, H. A. R. STAT activation and differential complex formation dictate selectivity of interferon responses. Acta Biochimica Polonica at 10.18388/abp.2007_3266 (2007).

41. Urbonaviciute, V., Luo, H., Sjöwall, C., Bengtsson, A. & Holmdahl, R. Low Production of Reactive Oxygen Species Drives Systemic Lupus Erythematosus. Trends in Molecular Medicine at 10.1016/j.molmed.2019.06.001 (2019).

42. Kelkka, T. et al. Reactive oxygen species deficiency induces autoimmunity with type 1 interferon signature. Antioxidants Redox Signal. (2014) doi:10.1089/ars.2013.5828.

43. Olsson, L. M. et al. A single nucleotide polymorphism in the NCF1 gene leading to reduced oxidative burst is associated with systemic lupus erythematosus. Ann. Rheum. Dis. (2017) doi:10.1136/annrheumdis-2017-211287.

44. Zhao, J. et al. A missense variant in NCF1 is associated with susceptibility to multiple autoimmune diseases. Nat. Genet. (2017) doi:10.1038/ng.3782.

45. Luo, H., et al. NCF1-dependent production of ROS protects against lupus by regulating plasmacytoid dendritic cell development and functions. JCI insight (2023).

46. Fitzgerald-Bocarsly, P., Dai, J. & Singh, S. Plasmacytoid dendritic cells and type I IFN: 50 years of convergent history. Cytokine Growth Factor Rev. (2008) doi:10.1016/j.cytogfr.2007.10.006.

47. Chen, Y. L. et al. A type I IFN-Flt3 ligand axis augments plasmacytoid dendritic cell development from common lymphoid progenitors. J. Exp. Med. (2013) doi:10.1084/jem.20130536.

48. Schlitzer, A. et al. Identification of CCR9- murine plasmacytoid DC precursors with plasticity to differentiate into conventional DCs. Blood (2011) doi:10.1182/blood-2010-12-326678.

49. Sareila, O., Jaakkola, N., Olofsson, P., Kelkka, T. & Holmdahl, R. Identification of a region in p47phox/NCF1 crucial for phagocytic NADPH oxidase (NOX2) activation. J. Leukoc. Biol. (2013) doi:10.1189/jlb.1211588.

50. Liu, S. et al. Phosphorylation of innate immune adaptor proteins MAVS, STING, and TRIF induces IRF3 activation. Science (80-.). (2015) doi:10.1126/science.aaa2630.

51. Weng, M.-T. & Luo, J. The enigmatic ERH protein: its role in cell cycle, RNA splicing and cancer. Protein Cell 4, 807–812 (2013).

52. Pang, K. et al. ERH gene and its role in cancer cells. Front. Oncol. 12, 900496 (2022).

53. Yu, Q. et al. Sample multiplexing for targeted pathway proteomics in aging mice. Proc. Natl. Acad. Sci. 117, 9723–9732 (2020).

54. O’Connell, J. D., Paulo, J. A., O’Brien, J. J. & Gygi, S. P. Proteome-wide evaluation of two common protein quantification methods. J. Proteome Res. 17, 1934–1942 (2018).

55. Végvári, Á., Rodriguez, J. E. & Zubarev, R. A. Single-cell chemical proteomics (SCCP) interrogates the timing and heterogeneity of cancer cell commitment to death. Anal. Chem. 94, 9261–9269 (2022).

56. Végvári, Á., Rodriguez, J. E. & Zubarev, R. A. Single Cell Proteomics Using Multiplexed Isobaric Labeling for Mass Spectrometric Analysis. Single-Cell Protein Anal. Methods Protoc. 113–127 (2022).

57. Mnatsakanyan, R. et al. Proteome-wide detection of S-nitrosylation targets and motifs using bioorthogonal cleavable-linker-based enrichment and switch technique. Nat. Commun. 10, 2195 (2019).

58. Zhang, X., Gaetani, M., Chernobrovkin, A. & Zubarev, R. A. Anticancer effect of deuterium depleted water - Redox disbalance leads to oxidative stress. Mol. Cell. Proteomics (2019) doi:10.1074/mcp.RA119.001455.

59. Gelderman, K. A. et al. Macrophages suppress T cell responses and arthritis development in mice by producing reactive oxygen species. J. Clin. Invest. 117, 3020–3028 (2007).

60. Szklarczyk, D. et al. The STRING database in 2017: quality-controlled protein–protein association networks, made broadly accessible. Nucleic Acids Res. gkw937 (2016).

61. Vizcaíno, J. A. et al. ProteomeXchange provides globally coordinated proteomics data submission and dissemination. Nature Biotechnology at 10.1038/nbt.2839 (2014).

