## Supplementary Information for "Multifaceted proteome analysis at solubility, redox, and expression dimensions for target identification"

**Supplementary Figures**

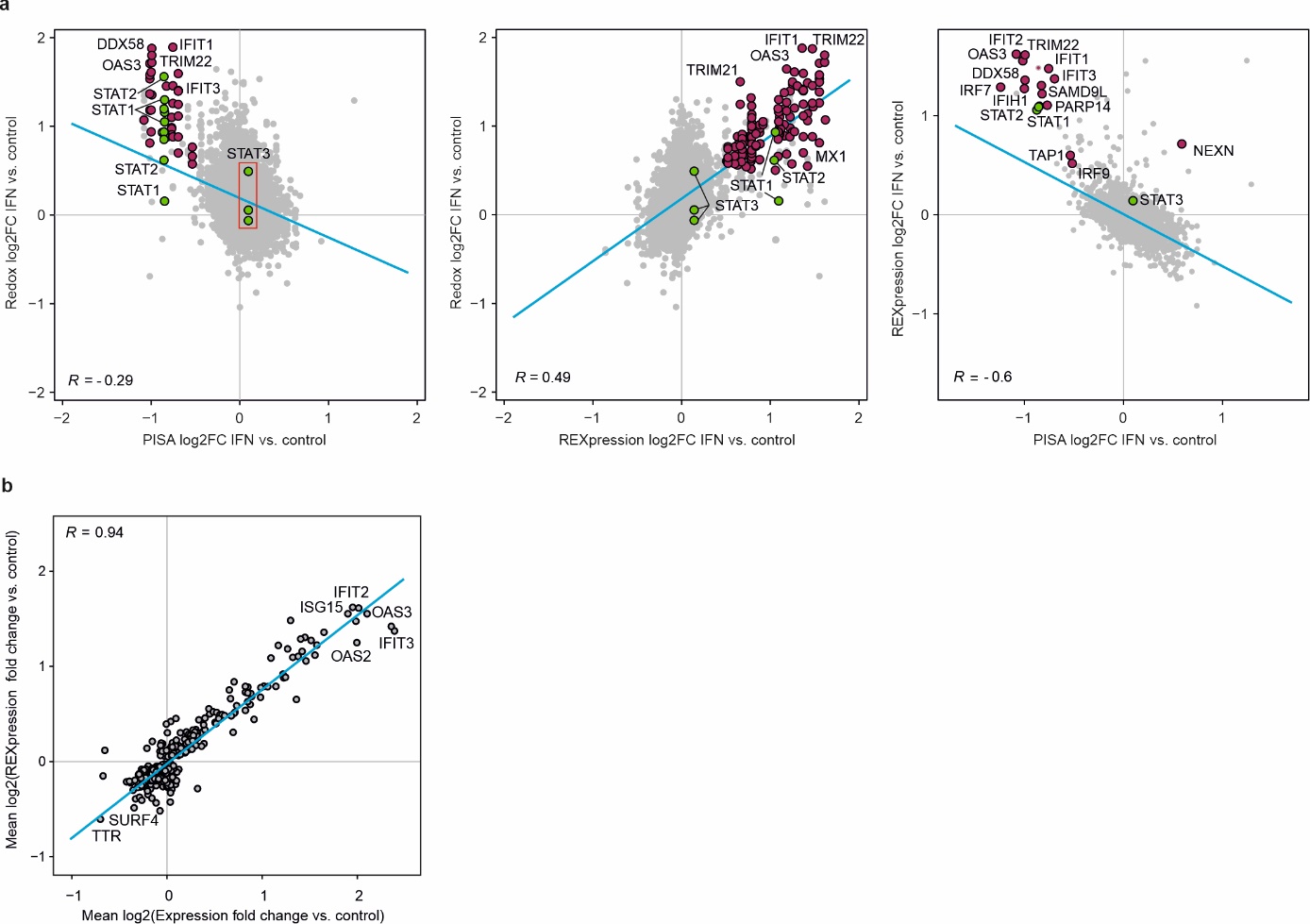

**Supplementary Fig. 1.** **a,** The scatterplots of three different dimensions highlight proteins changing across different dimensions upon IFN-α treatment. **b,** The correlation of protein fold changes upon IFN-α treatment vs. control, in REXpression vs. expression.

**Supplementary Tables**

**Supplementary** **Table 1.** The treatment concentrations and durations

| **Experiment** | **Cells and concentrations** | **PISA treat time** | **Expression treat time** | **REX treat time** |
| --- | --- | --- | --- | --- |
| Auranofin #1 | PISA and REX at 3 μM and expression at 1.5 μM | 2 h | 48 h | 2 h |
| Auranofin #2 | HCT116 (1.5 μM) | 24 h | 24 h | 24 h |
| Interferons α | THP1 (10 ng/mL) | 16 h | 16 h | 16 h |
| Interferons α | PDC cells from mutant and wt mice (500U/ml) | 20 h | 20 h | 20 h |

**Supplementary Table 2. Experimental design for the PISA-REX experiments.** the columns correspond to the TMT labels 1-16

| **1** | **2** | **3** | **4** | **5** | **6** | **7** | **8** | **9** | **10** | **11** | **12** | **13** | **14** | **15** | **16** |
| --- | --- | --- | --- | --- | --- | --- | --- | --- | --- | --- | --- | --- | --- | --- | --- |
| PISA control | PISA control | PISA treatment | PISA treatment | Expression control | Expression control | Expression control | Expression treatment | Expression treatment | Expression treatment | REX control | REX control | REX control | REX treatment | REX treatment | REX treatment |

**Supplementary Table 3.** Details of the LC-MS settings for the proteomic experiments performed in the current paper

| **Parameters/instrument** | **HF** | **Exploris** | **Lumos** |
| --- | --- | --- | --- |
| **Gradient time (min)** | 90 | 90 | 95 |
| **MS scan range (m/z)** | 375-1500 | 375-1500 | 375-1500 |
| **HCD collision energy** | 33 | 33 | 35 |
| **Orbitrap resolution** | 120,000 | 120,000 | 120,000 |
| **MS^2^ resolution** | 45,000 | 45,000 | 60,000 |
| **MS AGC target** | 3e6 | 3e6 | 2.5e5 (250%) |
| **MS^2^ AGC target** | 2e5 | 2e5 | 2.5e5 |
| **MS maximum injection time (ms)** | 100 | 50 | Auto |
| **MS^2^ maximum injection time**  **(ms)** | 120 | 120 | Auto |
| **Isolation window** | 1.6 | 1.6 | 1.6 |
